# Transcutaneous electrical nerve stimulation acutely impacts motor unit firing activity during isometric contractions

**DOI:** 10.1101/2022.12.22.521271

**Authors:** Simon Avrillon, Julio C. Hernandez-Pavon, Nish M. Kurukuti, Grace W. Hoo, José L. Pons

**Author notes:** **Correspondence:** Simon Avrillon, Imperial College London, London, UK, **e-mail:**, José L. Pons, Legs + Walking Lab, floor 23, Shirley Ryan AbilityLab, Chicago, IL, USA, **e-mail:**.

## Abstract

A low intensity electrical current delivered transcutaneously at a high frequency over a muscle can acutely recruit motor units in a physiological order by activating peripheral sensory pathways. This method has been used in patients to reduce tremor or supplement motor function, leading to the development of therapies and products.

We aimed to better understand how the stimulation of the median nerve, the contralateral first dorsal interosseus muscle (FDI), and the combination of these two paradigms impact the motor unit activity from the FDI muscle. We identified and tracked the same motor units across the conditions and compared the electromyographic amplitude, motor unit discharge rates, and the degree of correlation between fast and slow oscillations of motor unit discharge rates.

We found that the stimulation of the FDI muscle can acutely increase the electromyographic amplitude of the homonymous muscle on the contralateral side (F = 20.4; p < 0.001) while the discharge rate of motor units did not differ between the control and the stimulation condition (F = 0.2; p = 0.806). We did not observe any significant effect of the stimulation on the ratio of pairs of motor units with a significant correlation, showing that the stimulation barely impacted the distribution of correlated inputs to the pool of motor units. We did not observe short-term effects of the stimulation once it was discontinued.
Overall, these results showed that the specific stimulation of peripheral sensory pathways can acutely impact motor unit firing activity without disturbing the neural control of force.

**New & Noteworthy:** We identified and tracked the same motor units across stimulation and control conditions using high-density electromyography. We found that the specific stimulation of peripherial sensory pathways can acutely impact motor unit firing activity, likely due to the recruitment of additional motor units. At the same time, the degree of correlation between fast and slow oscillations of motor unit discharge rates was stable, limiting the disturbance of the neural control of force.

## Introduction

The modulation of muscle force requires coherent and common oscillations of motor units discharge rates, particularly in the bandwidth of force fluctuations (1, 2). These common oscillations arise from supraspinal centers and spinal circuits and converge to spinal motor neurons, the ‘final common path’ that transduces neural inputs into mechanical outputs (3). Besides generating excitatory and inhibitory post-synaptic inputs that modulate motor neuron firing activity (4, 5), spinal circuits can also amplify or reduce descending oscillations in specific bandwidths by shifting the phase of their oscillations (6, 7). For example, Williams and Baker (7, 8) proposed that Renshaw cells could actively attenuate oscillations linked to physiological tremors, i.e., around 10 Hz, by oscillating in antiphase at the same frequencies. This has functional and practical implications, e.g., peripheral electrical stimulation of afferent sensory fibers in antiphase with tremorgenic inputs could also attenuate pathologic tremors in patients and improve their motor function (9, 10). Interestingly, the effects of the stimulation may even remain once the electrical stimuli are discontinued, which implies that short-term adaptation may occur at the spinal and supraspinal levels (10).

Peripheral transcutaneous electrical stimulation has been used in clinical settings to supplement the activation of motor units (11). In this way, the electrical current applied over a muscle through conductive pads activates the motor units positioned in the underlying tissue, regardless of their recruitment threshold (12). Unfortunately, such stimulations may generate fatigue and discomfort for patients. For these reasons, numerous studies have focused on the selective activation of peripheral sensory pathways to activate motor units in a physiological order at the spinal level (13, 14). As sensory fibers have a lower rheobase than motor fibers, a current delivered at an amplitude below the motor threshold can selectively activate them and generate inhibitory or excitatory post-synaptic potentials reaching pools of motor neurons through various spinal circuits (13, 14). For example, studies have found that the transcranial electrical nerve stimulation of the biceps brachii on one limb can increase the EMG amplitude of the homonymous muscle in the contralateral limb (15). Other studies have found that supplemented sensory feedback could also improve motor functions in patients with multiple sclerosis (16) or stroke (17). It is noteworthy that the effect of supplemented sensory feedback on motor unit firing activity was not uniform (18), likely due to non-homogeneous distribution of afferent inputs to motor neurons from the same pool (19) and a size effect of afferent inputs onto motor neuron firing activity (20). Whether this divergent modulation of motor neuron firing activity within the same pool impacts the motor unit firing activity, and as a consequence, the modulation of force remains unclear.

We hypothesized that supplementing peripheral sensory pathways with transcutaneous electrical stimulation would i) acutely modulate motor unit firing activity, but ii) limit the degree of correlation between motor units firing activities from the same pool. To this end, we asked 12 participants to perform isometric abductions of the index finger while we recorded high-density surface electromyographic (HDsEMG) signals over the first dorsal interosseous (FDI) muscle. We compared four conditions with 1) stimulation over the median nerve on the same side, 2) stimulation over the FDI on the contralateral side, 3) condition 1 and 2, and 4) no stimulation. We decomposed the HDsEMG signals into individual motor unit spike trains (21) and tracked the same motor units across the four conditions using the unique spatiotemporal properties of motor unit action potentials (22). We compared the discharge characteristics of the tracked motor units, the EMG amplitude, and the degree of correlation between motor units firing activities over three bandwidths, i.e., 0-5 Hz, 0-10 Hz, and 0-20 Hz.

## Methods

### Participants

12 right-handed individuals volunteered to participate in the study (mean ± SD; Age: 30 ± 10 years; Height: 174 ± 10 cm; Body mass: 77 ± 14 kg; 6 women). They had no history of brain lesions, no neurological disorders, and normal vision with or without correction. They provided an informed written consent form during the enrollment session. The institutional review board of Northwestern University reviewed and approved the study (IRB#: STU00211930).

### Experimental setup

The experimental session consisted of a series of isometric abductions of the right index finger against a force sensor (MLP-200, Transducer Techniques, CA, USA) with or without transcutaneous peripheral electrical stimulations. The participant sat in an instrumented chair with adjustable angles and height to maximize their comfort. We aligned the right hand and forearm with the center of the force sensor, and we securely strapped the right arm and forearm to the arm rest to prevent any movement. We asked the participant to extend and separate the index finger from the middle, ring, and little fingers to only produce abduction force with the index finger. The force signal was digitized at 2048 Hz using the HDsEMG acquisition system (Quattrocento; 384 channels for bioelectrical signals and 16 channels for auxiliary inputs, OT Bioelettronica, Italy).

After a standardized warm-up, which consisted of 10 contractions at 50 %, three contractions at 70 %, and three contractions at 80 % of the subjective maximal voluntary contraction (MVC), the participant performed three MVC for 10 s with 90 s of rest in between. We applied a moving-averaged window of 250 ms over the force signals and considered the maximal value obtained across the three MVC as the maximal force. Then, we asked participants to perform a series of trapezoidal isometric contractions with 10 s ramp-up and ramp-down, and a 60 s plateau set at 10 % of the MVC. A screen positioned in front of the participant displayed a real-time visual force feedback and a target that the participant tracked. The participant performed three submaximal contractions with 30 s of rest in between per stimulation condition and two submaximal contractions for the control condition, for a total of 11 contractions. The order of conditions was randomized.

### Stimulation paradigms

We compared three stimulation paradigms with a control condition (Fig. 1A). Each stimulation condition aimed at supplementing peripheral sensory pathways during submaximal isometric contractions to modulate motor unit firing activity. We randomized three stimulation paradigms: i) stimulation of the right median nerve, ii) stimulation of the left FDI muscle belly, iii) stimulation of both the right median nerve and the left FDI muscle belly (Fig. 1B). Each stimulation pattern was delivered during three blocks of 30 s, for a total of nine blocks of 30 s during the entire session. Previous work showed that the activation of the median nerve, mostly composed of sensory fibers (23), impacted the activation of the FDI muscle (24). Similarly, transcutaneous electrical nerve stimulation of a muscle increased motor unit firing activity of the same muscle on the contralateral side (15).

**Figure 1.**
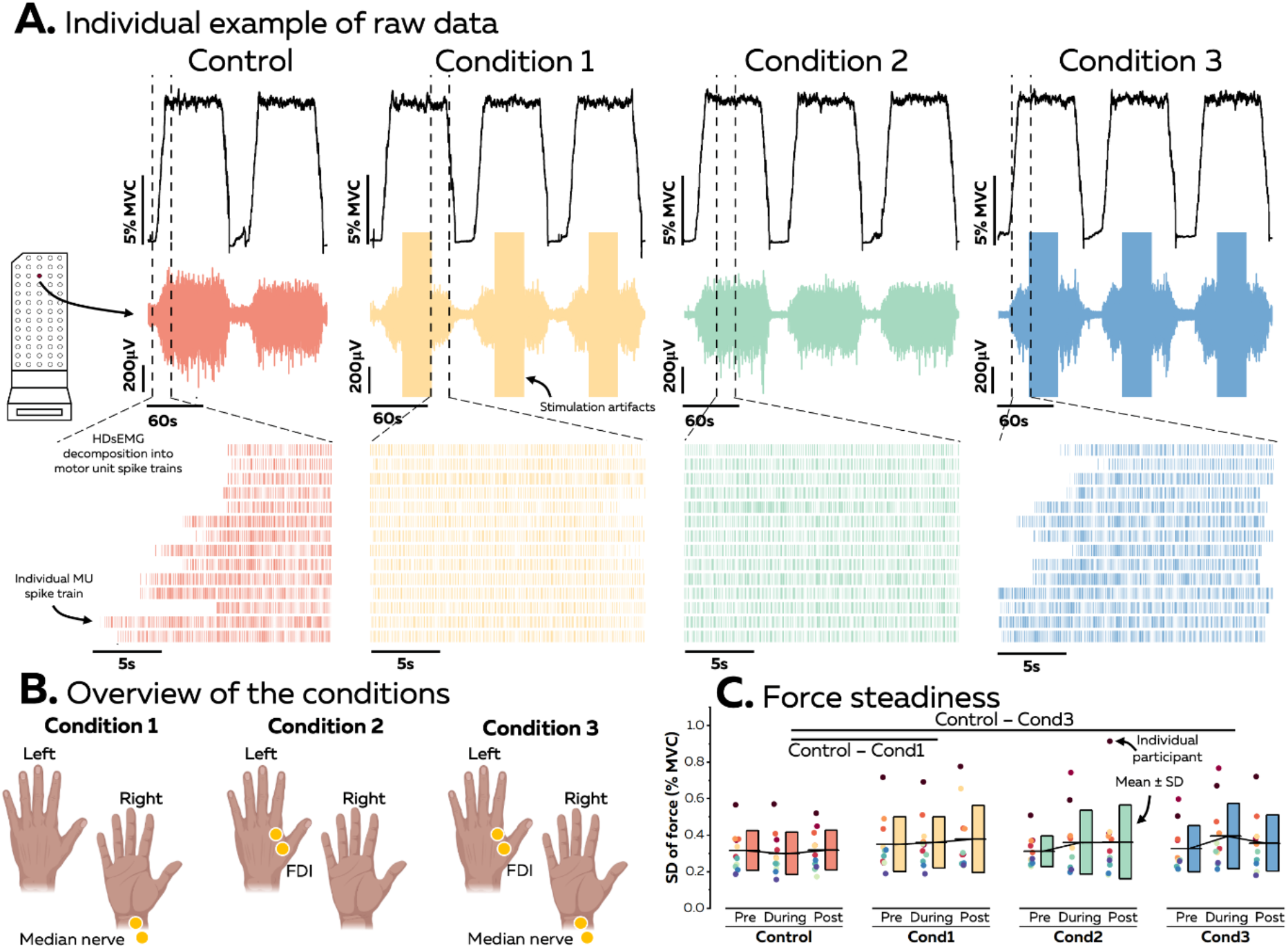
Overview of the data collection. (A) Each participant performed a series of trapezoidal isometric contractions at 10 % MVC during which we recorded high-density electromyographic signals over the FDI muscle. We decomposed these signals into individual motor unit spike trains and compared motor unit firing activity in four conditions. (B) Control: no stimulation; Condition 1: stimulation of the right median nerve; Condition 2: stimulation of the left FDI muscle belly; Condition 3: stimulation of both the median nerve and the left FDI muscle belly. (C) We first estimated force steadiness as the standard deviation of force during the plateau before, during, and after the stimulation. Each scatter represents individual data. The line is the mean. The boxes display ± one standard deviation. The horizontal lines denote a statistical difference between conditions.

We first used handheld dry electrodes with an anode and a cathode of 12 mm diameter separated by 50 mm and connected to a bipolar constant current stimulator (Digitimer DS5, Digitimer, UK) to find the optimal location that selectively activate the muscles of the thenar eminence - innervated by the median nerve - on the right side, and the FDI muscle on the left side. Then, the skin was cleaned with alcohol, and two round conductive pads were placed over the skin (PALS electrodes with a diameter of 2.5 cm, Axelgaard, USA). We then found the sensory threshold, i.e., the minimal current where the participant could feel the stimulation, and the motor threshold, i.e., the minimal current intensity where the muscle contraction was visible. The level of intensity was set to the highest current intensity below the motor threshold. When this current intensity was too painful for the participant, we lowered the current intensity until reaching a tolerable intensity, with a minimal intensity set to 1.5 × the sensory threshold. During the session, the stimulation was delivered at a frequency of 100 Hz with a pulse duration of 400 μs.

### HDsEMG recordings

HDsEMG signals were recorded from the right FDI muscle using a two-dimensional adhesive grid of 64 electrodes (13×5 electrodes with one electrode absent on a corner, gold-coated, inter-electrode distance: 4 mm; [GR04MM1305, OT Bioelettronica, Italy]). Before electrode application, the skin was shaved if necessary, and then cleaned with an abrasive pad and water. The adhesive grid was held on the skin using semi-disposable bi-adhesive foam layers (SpesMedica, Battipaglia, Italy). Skin-electrode contact was assured by filling the cavities of the adhesive layers with conductive paste. Strap electrodes dampened with water were placed around the contralateral (ground electrode) and ipsilateral (reference electrode) wrists. The EMG signals were recorded in monopolar mode, bandpass filtered (10-500 Hz) and digitized at a sampling rate of 2048 Hz using a multichannel acquisition system (EMG-Quattrocento; 400-channel EMG amplifier, OT Bioelettronica, Italy).

### HDsEMG decomposition

We decomposed HDsEMG signals before and after the stimulation for conditions 1 and 3, as the signals were contaminated by stimulation artifacts (Fig. 1A), and over the entire contraction for condition 2 and the control condition. First, the monopolar EMG signals were bandpass filtered between 20-500 Hz with a second-order Butterworth filter. After visual inspection, channels with low signal-to-noise ratio or artifacts were discarded. The HDsEMG signals were then decomposed into motor unit spike trains using convolutive blind-source separation (21). In short, the EMG signals were first extended and whitened. Thereafter, a fixed-point algorithm that maximized the sparsity was applied to identify the sources embedded in the EMG signals, i.e., the motor unit spike trains. Motor unit spike trains can be considered as sparse sources with most samples being 0 (i.e., absence of spikes) and a few samples being 1 (i.e., spikes). In this algorithm, a contrast function was iteratively applied to the EMG signals to estimate the level of sparsity of the identified source, and the convergence was reached once the level of sparsity did not vary when compared to the previous iteration, with a tolerance fixed at 10-4 (see (21) for the definition of the detailed contrast functions). At this stage, the estimated source contained high peaks (i.e., the spikes from the identified motor unit) and low peaks from other motor units and noise. High peaks were separated from low peaks and noise using peak detection and K-mean classification with two classes. The peaks from the class with the highest centroid were considered as the spikes of the identified motor unit. A second algorithm refined the estimation of the discharge times by iteratively recalculating the motor unit filter and repeating the steps with peak detection and K-mean classification until the coefficient of variation of the inter-spike intervals was minimized. This decomposition procedure has been previously validated using experimental and simulated signals (21). After the automatic identification of the motor units, duplicates were removed, and all the motor unit spike trains were visually checked for false positives and false negatives. This manual step is highly reliable across operators (25). As usually done, only the motor units which exhibited a pulse-to-noise ratio > 30 dB were retained for further analysis. This threshold ensured a sensitivity higher than 90 % and a false-alarm rate lower than 2 % (26).

### Motor unit tracking

To compare motor unit discharge characteristics across conditions, we needed to track the same motor units across separated decompositions. Therefore, we matched motor units using the unique spatio-temporal properties of motor unit action potentials within the grids of electrodes (Fig. 2A; (22, 27)). Specifically, the motor unit action potential shapes were identified using the spike-triggered averaging technique. As the position of the action potentials within the window may differ between motor units, we considered the local maxima, i.e., the action potential with the highest peak-to-peak amplitude within the grid, as the center of the 50 ms window. Finally, we concatenated the motor unit action potentials from the 64 channels and performed a 2D cross-correlation between motor units. Pairs with a correlation coefficient higher than 0.75 were considered as matches (22, 27). The matches were visually checked to guarantee the similarity of the motor unit action potential shapes. Only motor units tracked across the four conditions were retained for the analysis.

**Figure 2.**
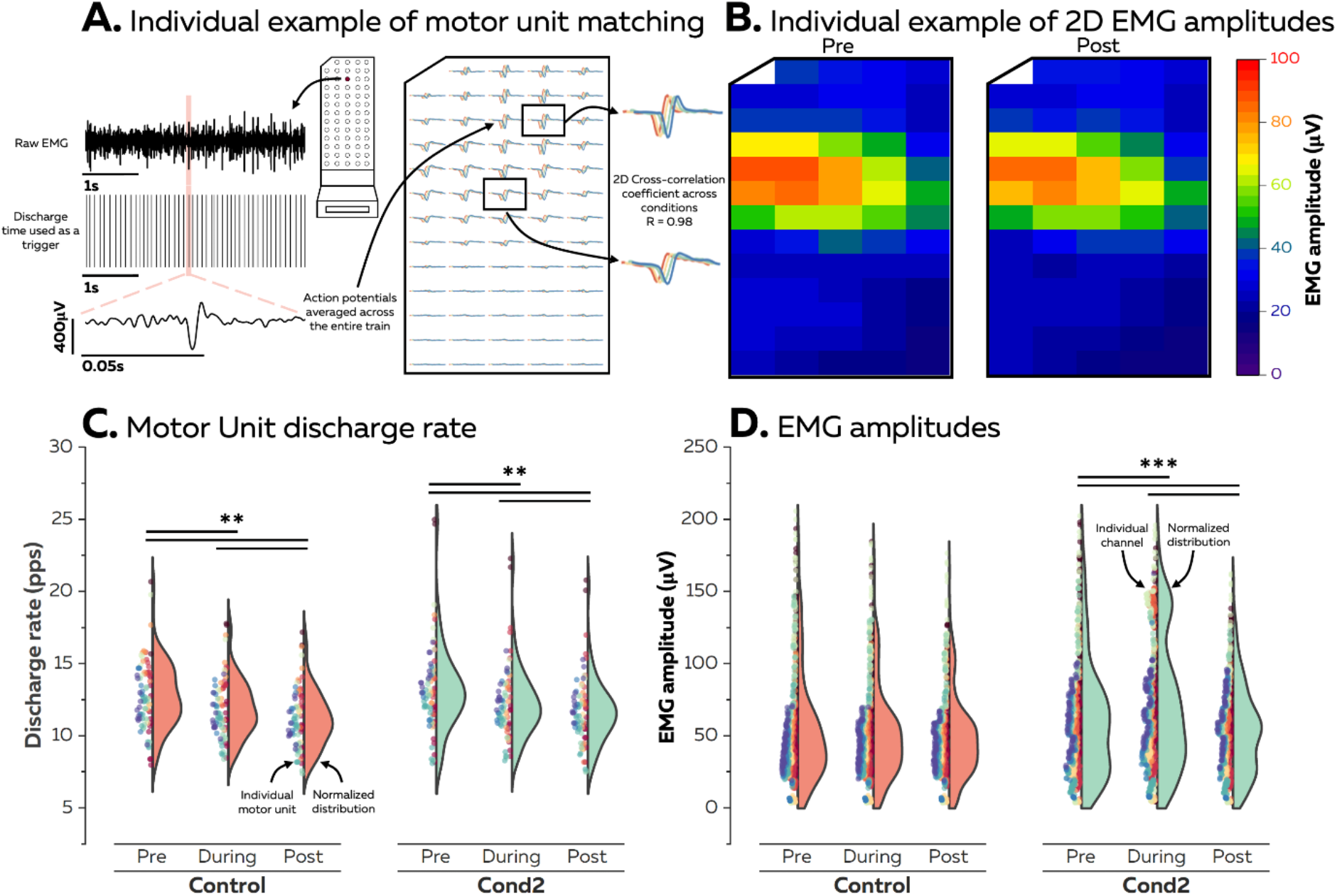
Acute changes in motor unit firing activity. (A) To compare the firing activity across the four conditions, we tracked the same motor units using 2D cross-correlation of the concatenated action potential waveforms. To this end, we used the discharge times as triggers to segment the electromyographic signal over each channel. We then averaged these waveforms over the entire spike trains and concatenated the waveforms from the 64 channels. 2D cross-correlations were performed for each pair of motor units, and pairs with a correlation coefficient higher than 0.75 were considered as matches. (B) We calculated the EMG amplitude before, during, and after the stimulation over the 64 channels by calculating averaged-rectified values. (C) Changes in discharge rates before, during, and after the stimulation are reported for all the matched motor unit. Each scatter is an individual motor unit with one color per participant. (D) Changes in EMG amplitude before, during, and after the stimulation are reported for all the individual channels. Each scatter is an individual channel with one color per participant. The horizontal lines denote a statistical difference between times within each condition.

### EMG amplitude and motor unit discharge characteristics

The monopolar EMG signals were bandpass filtered between 10 and 500 Hz with a third-order Butterworth filter. Each channel was visually inspected and channels with low signal-to-noise ratio or artifacts were discarded. Each signal was rectified and averaged over non-overlapping windows of 1s. We estimated the EMG amplitude for each channel as the mean of averaged-rectified values before, during, and after the transcutaneous electrical stimulation, except for conditions 1 and 3 where stimulation artifacts prevented us to estimate EMG amplitude during the stimulation (Fig. 2B). Instantaneous motor unit discharge rates were calculated and averaged over the same windows.

### Correlation between motor units firing activities

We estimated the correlation between motor unit smoothed discharge rates over various bandwidths. To this end, the decomposed motor unit spike times were first converted into continuous binary signals with ones corresponding to the firing instances of a unit. The smoothed discharge rates were then obtained by convoluting these binary signals with 200 ms, 100 ms, and 50 ms Hanning windows. This respectively corresponds to bandwidths of 0-5 Hz, 0-10 Hz, and 0-20 Hz. These signals were high pass filtered with a cut-off frequency of 0.75 Hz to remove offset and trends (28). We calculated the cross-correlation coefficients for each pair of motor units on their smoothed discharge rates. The maximal cross-correlation coefficients within a time lag of −100 to +100 ms were considered for analysis.

Besides correlation amplitudes, we also reported the number of pairs of motor units with statistically significant correlation coefficients (29, 30). Thus, we considered that the pairs of motor units with significant correlation coefficients received correlated inputs. To this end, we defined a significance threshold as the 95th percentile of the cross-correlation coefficient distribution generated with resampled versions of all motor unit spike trains per participant and condition. We then generated random spike trains for each motor unit by bootstrapping the interspike intervals (random sampling with replacement). This random spike train had the same number of spikes and the same discharge rate (mean and standard deviation) as the original motor unit spike train. We repeated this step four times per pair of motor units.

### Statistics

All statistical analyses were performed with RStudio (Boston, MA, USA). First, quantile-quantile plots and histograms were displayed to check the normality of the data distribution. If the data were determined not to be normal, they were transformed to remove the skew. The first analyses were performed using linear mixed effect models implemented in the R package lmerTest with the Kenward-Roger method to estimate the denominator degrees of freedom and the p-values. This method considers the dependence of data points within each participant due to, i.e., common synaptic inputs that impact the behavior of motor units from the same pool. When necessary, multiple comparisons were performed using the R package emmeans, which adjusts the p-value using the Tukey method. The significance level was set at p ≤ 0.05. Values are reported as mean ± standard deviation.

To compare the force steadiness between conditions, we calculated the standard deviation of force and used a linear mixed effects model with Time (before, during, after) and Condition (Control, Stimulation #1, Stimulation #2, Stimulation #3) as fixed effects and Participants as a random effect.

To assess the acute effect of transcutaneous electrical stimulation on EMG amplitude and motor unit discharge rates, we used a linear mixed effects model with Time (before, during, after) and Condition (Control, Stimulation #2) as fixed effects and Participants as a random effect. To assess the acute effect of transcutaneous electrical stimulation on the degree of correlation between motor unit smoothed discharge rates over 0-5 Hz, 0-10 Hz, and 0-20 Hz, we used a linear mixed effects model for each bandwidth with Time (before, during, after) and Condition (Control, Stimulation #2) as fixed effects and Participants as a random effect.

To assess the short-term effect of transcutaneous electrical stimulation on EMG amplitude and motor unit discharge rates, we used a linear mixed effects model with Time (before, during, after) and Condition (Control, Stimulation #1, Stimulation #2, Stimulation #3) as fixed effects and Participants as a random effect. To assess the short-term effect of transcutaneous electrical stimulation on the degree of correlation between motor unit smoothed discharge rates over 0-5 Hz, 0-10 Hz, and 0-20 Hz, we used a linear mixed effects model for each bandwidth with Time (before, during, after) and Condition (Control, Stimulation #1, Stimulation #2, Stimulation #3) as fixed effects and Participants as a random effect.

## Results

All the participants successfully tracked the target, with a force over the plateau of 10.2 ± 0.3 % MVC, 9.9 ± 0.3 % MVC, 9.8 ± 0.6 % MVC, and 10.0 ± 0.3 % MVC for Control, Conditions 1, 2, and 3, respectively. We then estimated the force steadiness by calculating the standard deviation of the force over the plateau before, during, and after the stimulation (or no stimulation for the control condition; Fig. 1C). We found a significant effect of Condition (F = 3.4; p = 0.020), but no effect of Time (F = 1.7; p = 0.194) nor an interaction between Time and Condition (F = 0.8; p = 0.579). Specifically, the standard deviation of the force was lower for Control (0.31 ± 0.11) than Condition 1 (0.36 ± 0.15; p = 0.024) and Condition 3 (0.36 ± 0.15; p = 0.047). We then decomposed the HDsEMG signals recorded over the FDI muscle to identify individual motor unit spike trains. On average, we identified 12 ± 3, 13 ± 4, 14 ± 4, and 13 ± 4 motor units for Control, Conditions 1, 2, and 3, respectively. We matched 8 ± 3 motor units (62.8 ± 13.9 % of identified motor units) per participant.

We first investigated the acute effect of stimulation on motor unit discharge characteristics. It is noteworthy that this analysis was done by comparing Control with Condition 2, as stimulation artifacts contaminated signals recorded during Conditions 1 and 3. When considering motor unit discharge rates, we found significant effects of Condition (F = 8.7; p = 0.003) and Time (F = 40.1; p < 0.001) while the interaction between Time and Condition was not significant (F = 0.2; p = 0.806; Fig. 2C). Specifically, the discharge rate constantly decreased over the plateau for both conditions and was on average lower for Control (12.0 ± 2.2 pps) comparing to Condition 2 (12.4 ± 2.6 pps; p = 0.003). When considering the EMG amplitude, we observed significant effects of Condition (F = 97.1; p < 0.001) and Time (F = 32.9; p < 0.001) with a significant interaction between Condition and Time (F = 20.4; p < 0.001; Fig. 2D). Specifically, EMG amplitude only increased during the Condition 2, with higher EMG amplitudes during the stimulation (72 ± 46 μV) comparing to before (62 ± 41 μV; p < 0.001) and after (54 ± 32; p < 0.001) the stimulation. No differences were observed in the Control condition, with EMG amplitudes of 56 ± 40 μV, 53 ± 34 μV, and 50 ± 30 μV at the beginning, middle, and end of the plateau, respectively.

We then looked at the degree of correlation between motor unit smoothed discharge rates over three bandwidths: 0-5 Hz, 0-10 Hz, and 0-20 Hz (Fig. 3). This analysis was performed on 240 pairs of matched motor units. When considering the bandwidth 0-5 Hz, we found main effects of Condition (F = 30.0; p <0.001) and Time (F = 33.7; p < 0.001) with a significant interaction between Time and Condition (F = 4.0; p = 0.019). The degree of correlation increased for both conditions over the middle of the plateau, with the stimulation ‘off and ‘on’ for Control and Condition 2, respectively. However, we did not observe any significant differences between Conditions (F = 1.3; p = 0.265) and Times (F = 0.5; p = 0.622) for the ratios of significant correlations (Fig. 3, bottom panels). On average, 92.6 ± 10.2 % of the pairs of motor units for the Control condition and 89.4 ± 11.0 % of the pairs or motor units for Condition 2 had significant correlations. This means that most of pairs of motor units received a significant degree of common inputs over the entire plateau for each condition. When considering the bandwidth 0-10 Hz, we found main effects of Condition (F = 12.8; p < 0.001) and Time (F = 6.9; p < 0.001), but no significant interaction between Time and Condition (F = 4.0; p = 0.019). There was also a significant effect of Condition (F = 5.8; p = 0.021) for the ratios of significant correlations, with 55.5 ± 23.1 % of the pairs of motor units for the Control and 46.2 ± 11.5 % of the pairs or motor units for Condition 2 with significant correlations. Finally, when considering the bandwidth 0-20 Hz, we found a main effect of Condition (F = 12.9; p < 0.001), but no effect of Time (F = 2.5; p = 0.081) nor a significant interaction between Time and Condition (F = 0.3; p = 0.770). No significant differences were observed between the ratios of significant correlations.

**Figure 3.**
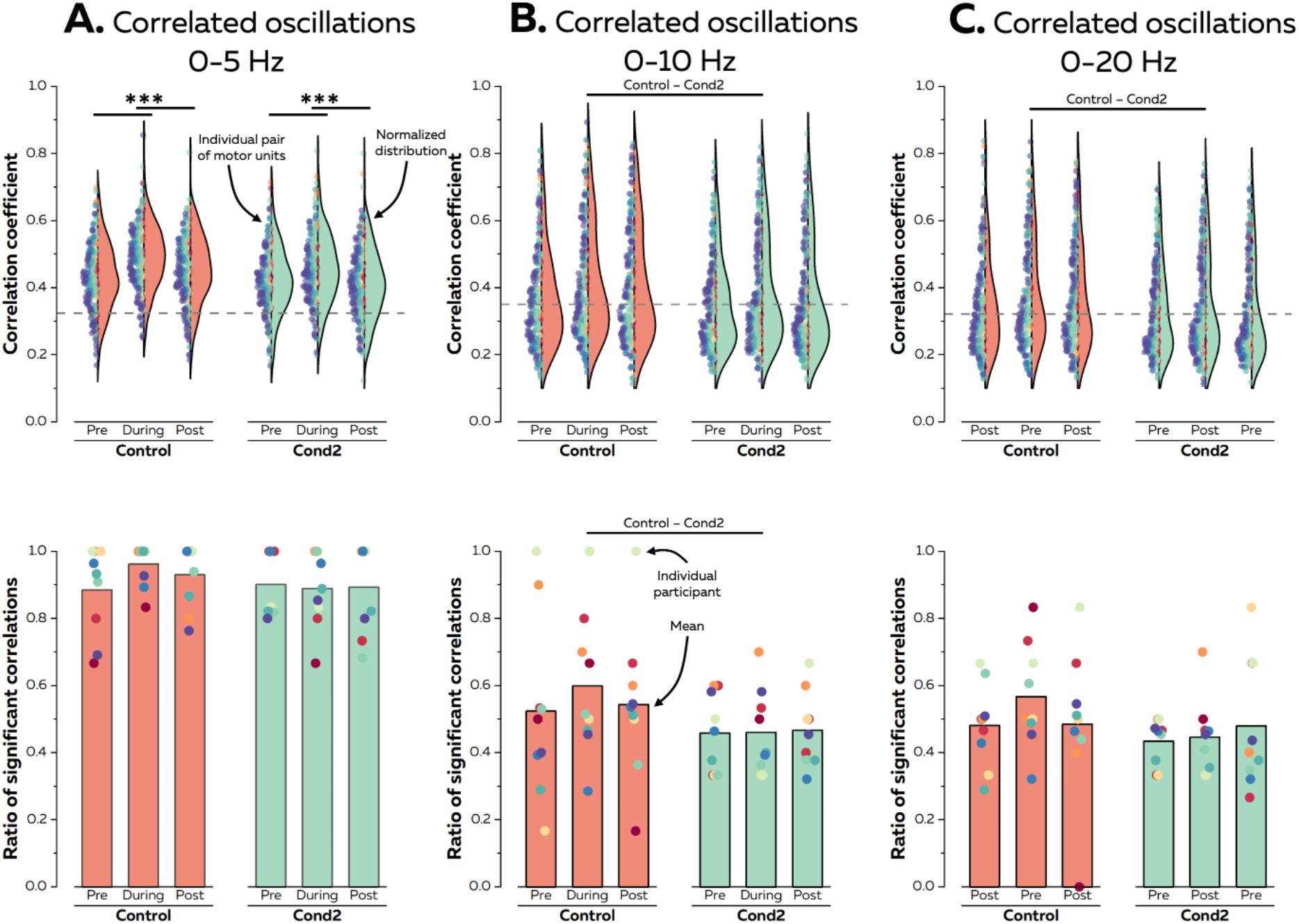
Acute changes in the level of correlation between motor unit firing activities. We estimated the correlation between motor unit smoothed discharge rates over 0-5 Hz (A), 0-10 Hz (B), and 0-20 Hz (C) bandwidths. For each analysis, we convoluted the motor unit spike trains with 200 ms, 100 ms, and 50 ms Hanning windows. This respectively corresponds to bandwidths of 0-5 Hz, 0-10 Hz, and 0-20 Hz. We reported the changes between correlation coefficients for each pair of matched motor units and the ratio of significant correlations out of the total number of correlations. On the top plots, each scatter is an individual pair of motor units with one color per participant. On the bottom plots, each scatter is an individual participant, and the bar is the mean. The horizontal lines denote a statistical difference between conditions. For a sake of clarity, we only display here the Condition effect or the interaction between Condition and Time.

Finally, we assessed the short-term effect of the stimulation on motor unit firing activity once the stimulation is discontinued. We performed this analysis on the three stimulation and control conditions (Fig. 4). We first compared motor unit discharge characteristics before and after the stimulation. When considering motor unit discharge rates, we found main effects of Condition (F = 5.2; p = 0.001) and Time (F = 157.6; p < 0.001), but no interaction between Condition and Time (F = 0.1; p = 0.949; Fig. 4A). When considering EMG amplitudes, we found main effects of Condition (F = 10.1; p < 0.001) and Time (F = 43.4; p < 0.001), but no interaction between Condition and Time (F = 0.4; p = 0.751; Fig. 4B). These two results showed that the stimulation did not impact the changes in motor unit discharge characteristics during the isometric contractions, with the same trends for the control and the three stimulation conditions.

**Figure 4.**
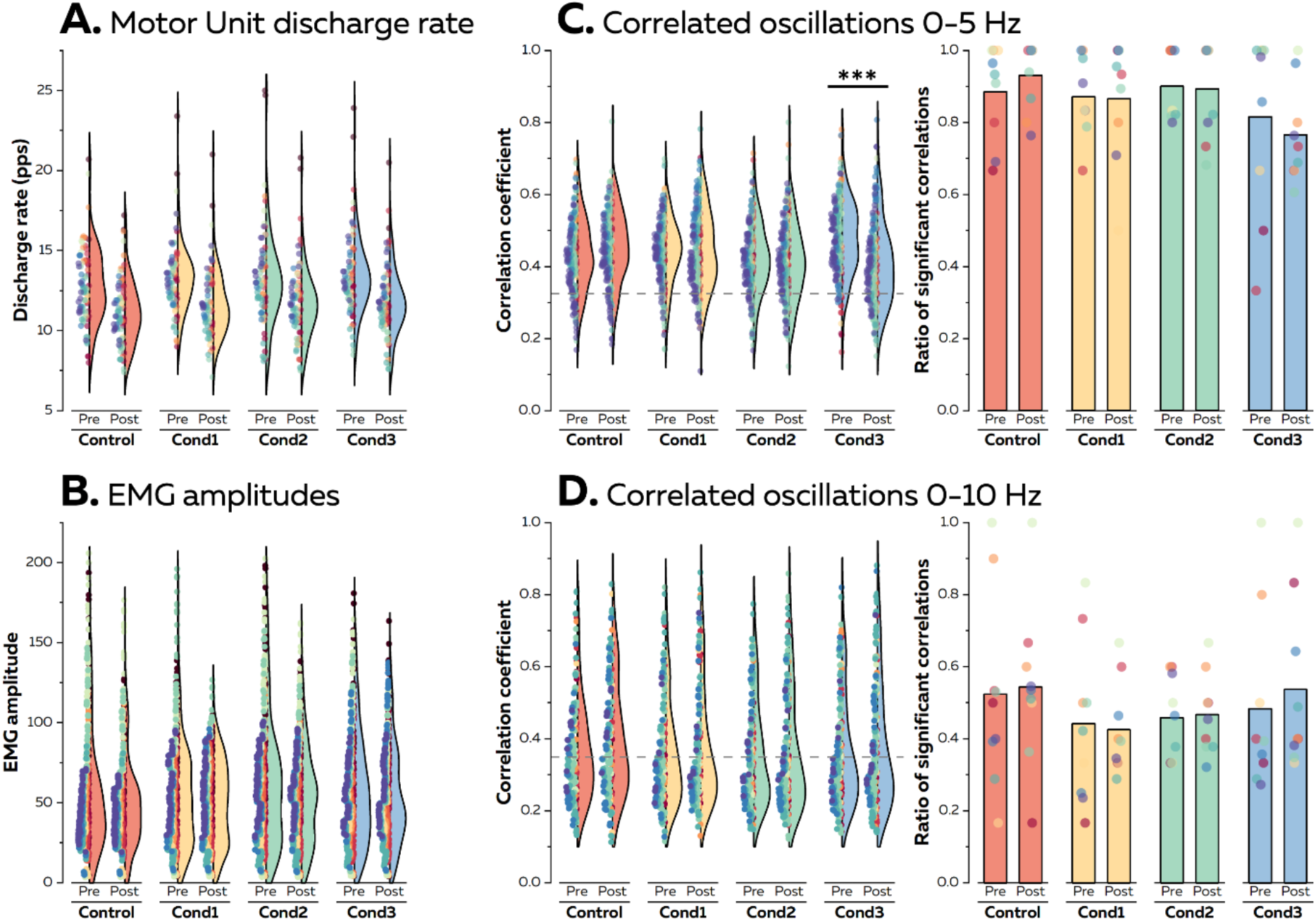
Short-term changes in motor unit firing activity. (A) We compared the discharge rate of each matched motor unit before and after the stimulation. Each scatter is an individual motor unit with one color per participant. (B) We compared the EMG amplitude of each channel before and after the stimulation. Each scatter is an individual channel with one color per participant. Note that it was not possible to estimate the discharge rate or to calculate EMG amplitude during the stimulation for Condition 1 and Condition 3 due to stimulation artifacts. (C) and (D), we compared the level of correlation between motor unit smooth discharge rates over two bandwidths: 0-5 Hz (C) and 0-10 Hz (D). We reported the differences in level of correlation between pairs of matched motor units and the ratio of significant correlations out of the total number of correlations. For the level of correlation, each scatter is an individual pair of motor units with one color per participant. Each scatter is an individual participant for the ratio of significant correlations, and the bar is the mean. The horizontal lines denote a statistical difference between times within each condition. For a sake of clarity, we only display here the interaction between Condition and Time

We finally assessed the degree of correlation between motor unit smoothed discharge rates over two bandwidths: 0-5 Hz and 0-10 Hz. For 0-5 Hz, we found main effects of Condition (F = 6.7; p < 0.001) and Time (F = 8.3; p = 0.003) with a significant interaction between Time and Condition (F = 19.2; p < 0.001). Specifically, the level of correlation between smoothed discharge rates decreased after the stimulation for Condition 3 (0.48 ± 0.10 vs. 0.41 ± 0.12; p < 0.001). However, we did not find any difference between Condition (F= 2.6; p = 0.063) or Time (F= 0.1; p = 0.891) for the ratios of significant correlation. This means that the number of pairs of motor units receiving significant common inputs did not change after the stimulation. For 0-10 Hz, we found main effects of Condition (F = 3.3; p = 0.019) and Time (F = 8.3; p = 0.004), but no significant interaction between Time and Condition (F = 0.5; p = 0.662). As for 0-5 Hz, we did not find any difference between Conditions (F= 1.7; p = 0.181) or Times (F= 0.2; p = 0.642) for the ratios of significant correlations.

## Discussion

In this study, we found that the specific stimulation of peripheral sensory pathways can acutely impact motor unit firing activity. Specifically, the EMG amplitude recorded over the FDI on the right side increased during the stimulation of the same muscle on the contralateral side. This change in EMG amplitude was likely due to the recruitment of additional motor units as the discharge rates of the identified motor units did not increase during the stimulation. Additionally, the number of pairs of motor units with significant correlations between their smoothed discharge rates did not differ between the Control and the Stimulation condition. This implies that the pool of motor units received a high degree of correlated inputs, which was barely impacted by the activation of afferent sensory fibers with electrical stimulation. Finally, we did not show any significant short-term effect of the stimulation on motor unit firing activity. Overall, these results confirm that the specific activation of peripheral sensory pathways can acutely supplement motor unit activity. However, contrary to our hypothesis, this does not acutely impact the degree of correlation between smoothed discharge rates and force steadiness.

The main result of our study is the acute effect of the specific activation of peripheral sensory pathways on motor unit activity. This is in line with previous studies showing that the stimulation of the biceps brachii acutely impacts motor unit activity of the same muscle on the contralateral side (15, 18), though our stimulation parameters (i.e., pulse width and frequency) were slightly different. The changes in motor unit activity are likely due to the generation of excitatory post-synaptic potentials at the spinal level, through several potential neural pathways. For example, several studies have shown that transcutaneous electrical nerve stimulation primarily activates afferent Ia fibers, thus recruiting the smallest motor units at the spinal level (31, 32). If single pulses below the motor threshold can elicit H-reflexes (33, 34), trains of stimulations can generate a sustained activation of these motor neurons until the stimulation is discontinued (31, 32). Others have found that the stimulation of cutaneous afferent fibers can also increase motor neuron excitability (35, 36). Importantly, the activation of these afferent sensory fibers can activate polysynaptic circuits across limbs (37), which explains why the stimulation of the FDI on the left side in this study elicited a response in the contralateral limb.

The changes in motor unit firing activity were likely due to the spatial recruitment of additional motor units as the discharge rates of the identified motor units did not change during the stimulation (Fig. 2D). This is again consistent with a previous study that observed the same adaptation when stimulating the contralateral limb (15). However, the fact that we observed a change in EMG amplitude without a change in motor unit discharge rate seems somewhat surprising as an increase in the synaptic input to the entire pool of motor neurons should also alter the discharge rate for the motor neurons already recruited. One explanation may be that the FDI muscle has several functional compartments where motor unit recruitment strategies appear to be independent of each other depending on the direction of the movement (38, 39). These divergent strategies are explained by the non-homogenous distribution of common inputs to motor neurons from the same muscle, with a higher degree of synchronization within compartments than between compartments (40, 41). Therefore, the increase in EMG amplitude may result from the recruitment of the smallest motor units from the non-activated compartments during the index finger’s abduction.

The second aim of the study was to assess the changes in the degree of correlation between smoothed motor unit discharge rates (28). To this end, we reported the level of correlation and the ratio of significant correlation coefficients. It is noteworthy that this method only estimates whether the pairs of motor units receive or do not receive correlated inputs (42), as the degree of correlation in the output is not linearly related to the degree of correlation between synaptic inputs (43). Contrary to our initial hypothesis, we did not find any differences between the Control and the Stimulation conditions over the same segment of the contraction. Note that in some cases, the ratio of significant correlations was lower for the stimulation condition over the entire contraction (Fig. 3). This implies that the supplemented peripheral sensory pathways did not significantly interfere with the overall level of correlated inputs. Spinal circuits may have filtered out these oscillations to prevent interference with force control (8). Indeed, previous work have shown that the stimulation of afferent fibers may increase physiological tremor (44, 45). Alternatively, this could potentially mean that the sensory contribution to the neural drive to motor unit was limited during our task, with most of the drive descending from supraspinal centers (46). However, it is difficult to transfer this finding to natural behaviors, as numerous studies have highlighted the key role of sensory feedback to regulate muscle coordination (47, 48) The impact of the specific activation of peripheral sensory pathways during movements remains to be determined in future studies. To this end, methods that identify and track individual motor units while the muscle geometry changes due to the movements will be required.

Finally, we did not find any short-term effect on the motor unit activity once the stimulation was discontinued (Fig. 4). We can therefore conclude that the effect of the stimulation was purely spinal and consisted of the generation of excitatory post-synaptic potentials that acutely affected motor neuron excitability. On the contrary, previous experiments have shown a ‘priming effect’ of such stimulations on motor neuron excitability, even when the stimulation was discontinued (36, 49). It is noteworthy that these studies used longer phases of stimulation (e.g., up to 40 min (49)). Other studies have found that the specific activation of sensory fibers may decrease tremor amplitude in patients with Essential Tremor after repeated phases of stimulation (10, 50). Practically, this means that longer stimulation sessions are needed to impact the excitability of the nervous system beyond the acute phase of the stimulation.

## Conclusion

The specific stimulation of peripheral sensory pathways acutely increased EMG amplitude while the discharge rate of motor units did not differ between the control and the stimulation conditions. It is likely that excitatory post-synaptic potentials were generated at the spinal level during the stimulation, thus recruiting the smallest motor units. Additionally, the degree of correlation between motor unit smoothed discharge rates was high and constant over the entire contraction, with or without stimulation. Therefore, we can conclude that the specific stimulation of peripheral sensory pathways can acutely impact motor unit firing activity without disturbing the neural control of force. As we did not observe short-term changes in motor unit firing activity once the stimulation was discontinued, we suggest that longer periods of stimulation should be used in a clinical setting to have an effect beyond the acute phase of stimulation.

## Acknowledgements

All the authors in this paper have no financial or other relationships that might lead to a conflict of interest.

## References

1. Farina D, Negro F, and Dideriksen JL. The effective neural drive to muscles is the common synaptic input to motor neurons. J Physiol 592: 3427–3441, 2014.

2. Enoka RM, and Farina D. Force Steadiness: From Motor Units to Voluntary Actions. Physiology (Bethesda) 36: 114–130, 2021.

3. Sherrington CS. Remarks on some aspects of reflex inhibition. Proc Biol Sci 97: 519–545, 1925.

4. Yavuz US, Negro F, Sebik O, Holobar A, Frommel C, Turker KS, and Farina D. Estimating reflex responses in large populations of motor units by decomposition of the high-density surface electromyogram. J Physiol 593: 4305–4318, 2015.

5. Nielsen JB. Human Spinal Motor Control. Annual review of neuroscience 39: 81–101, 2016.

6. Harel R, Asher I, Cohen O, Israel Z, Shalit U, Yanai Y, Zinger N, and Prut Y. Computation in spinal circuitry: lessons from behaving primates. Behav Brain Res 194: 119–128, 2008.

7. Williams ER, Soteropoulos DS, and Baker SN. Spinal interneuron circuits reduce approximately 10-Hz movement discontinuities by phase cancellation. Proc Natl Acad Sci U S A 107: 11098–11103, 2010.

8. Williams ER, and Baker SN. Renshaw cell recurrent inhibition improves physiological tremor by reducing corticomuscular coupling at 10 Hz. J Neurosci 29: 6616–6624, 2009.

9. Dideriksen JL, Laine CM, Dosen S, Muceli S, Rocon E, Pons JL, Benito-Leon J, and Farina D. Electrical Stimulation of Afferent Pathways for the Suppression of Pathological Tremor. Front Neurosci 11: 178, 2017.

10. Pascual-Valdunciel A, Gonzalez-Sanchez M, Muceli S, Adan-Barrientos B, Escobar-Segura V, Perez-Sanchez JR, Jung MK, Schneider A, Hoffmann KP, Moreno JC, Grandas F, Farina D, Pons JL, and Barroso FO. Intramuscular Stimulation of Muscle Afferents Attains Prolonged Tremor Reduction in Essential Tremor Patients. IEEE Trans Biomed Eng 68: 1768–1776, 2021.

11. Enoka RM, Amiridis IG, and Duchateau J. Electrical Stimulation of Muscle: Electrophysiology and Rehabilitation. Physiology (Bethesda) 35: 40–56, 2020.

12. Feiereisen P, Duchateau J, and Hainaut K. Motor unit recruitment order during voluntary and electrically induced contractions in the tibialis anterior. Exp Brain Res 114: 117–123, 1997.

13. Collins DF. Central contributions to contractions evoked by tetanic neuromuscular electrical stimulation. Exerc Sport Sci Rev 35: 102–109, 2007.

14. Dideriksen JL, Muceli S, Dosen S, Laine CM, and Farina D. Physiological recruitment of motor units by high-frequency electrical stimulation of afferent pathways. J Appl Physiol (1985) 118: 365–376, 2015.

15. Hamilton LD, Mani D, Almuklass AM, Davis LA, Vieira T, Botter A, and Enoka RM. Electrical nerve stimulation modulates motor unit activity in contralateral biceps brachii during steady isometric contractions. J Neurophysiol 120: 2603–2613, 2018.

16. Almuklass AM, Capobianco RA, Feeney DF, Alvarez E, and Enoka RM. Sensory nerve stimulation causes an immediate improvement in motor function of persons with multiple sclerosis: A pilot study. Mult Scler Relat Disord 38: 101508, 2020.

17. Celnik P, Hummel F, Harris-Love M, Wolk R, and Cohen LG. Somatosensory stimulation enhances the effects of training functional hand tasks in patients with chronic stroke. Arch Phys Med Rehabil 88: 1369–1376, 2007.

18. Amiridis IG, Mani D, Almuklass A, Matkowski B, Gould JR, and Enoka RM. Modulation of motor unit activity in biceps brachii by neuromuscular electrical stimulation applied to the contralateral arm. J Appl Physiol (1985) 118: 1544–1552, 2015.

19. Powers RK, and Binder MD. Distribution of oligosynaptic group I input to the cat medial gastrocnemius motoneuron pool. J Neurophysiol 53: 497–517, 1985.

20. Bawa P, Binder MD, Ruenzel P, and Henneman E. Recruitment order of motoneurons in stretch reflexes is highly correlated with their axonal conduction velocity. J Neurophysiol 52: 410–420, 1984.

21. Negro F, Muceli S, Castronovo AM, Holobar A, and Farina D. Multi-channel intramuscular and surface EMG decomposition by convolutive blind source separation. J Neural Eng 13: 026027, 2016.

22. Martinez-Valdes E, Negro F, Laine CM, Falla D, Mayer F, and Farina D. Tracking motor units longitudinally across experimental sessions with high-density surface electromyography. J Physiol 595: 1479–1496, 2017.

23. Gesslbauer B, Hruby LA, Roche AD, Farina D, Blumer R, and Aszmann OC. Axonal components of nerves innervating the human arm. Ann Neurol 2017.

24. Rossini PM, Rossi S, Tecchio F, Pasqualetti P, Finazzi-Agro A, and Sabato A. Focal brain stimulation in healthy humans: motor maps changes following partial hand sensory deprivation. Neurosci Lett 214: 191–195, 1996.

25. Hug F, Avrillon S, Del Vecchio A, Casolo A, Ibanez J, Nuccio S, Rossato J, Holobar A, and Farina D. Analysis of motor unit spike trains estimated from high-density surface electromyography is highly reliable across operators. J Electromyogr Kinesiol 58: 102548, 2021.

26. Holobar A, Minetto MA, and Farina D. Accurate identification of motor unit discharge patterns from high-density surface EMG and validation with a novel signal-based performance metric. J Neural Eng 11: 016008, 2014.

27. Vecchio AD, and Farina D. Interfacing the neural output of the spinal cord: robust and reliable longitudinal identification of motor neurons in humans. J Neural Eng 17: 016003, 2019.

28. De Luca CJ, and Erim Z. Common drive in motor units of a synergistic muscle pair. J Neurophysiol 87: 2200–2204, 2002.

29. Hug F, Avrillon S, Sarcher A, Del Vecchio A, and Farina D. Correlation networks of spinal motor neurons that innervate lower limb muscles during a multi-joint isometric task. J Physiol 2022.

30. Maillet J, Avrillon S, Nordez A, Rossi J, and Hug F. Handedness is associated with less common input to spinal motor neurons innervating different hand muscles. J Neurophysiol 128: 778–789, 2022.

31. Blouin JS, Walsh LD, Nickolls P, and Gandevia SC. High-frequency submaximal stimulation over muscle evokes centrally generated forces in human upper limb skeletal muscles. J Appl Physiol (1985) 106: 370–377, 2009.

32. Collins DF, Burke D, and Gandevia SC. Sustained contractions produced by plateau-like behaviour in human motoneurones. J Physiol 538: 289–301, 2002.

33. Hoffmann P. Ueber die Beziehungen der Sehnenreflexe zur willkürlichen Bewegung und zum Tonus. R. Oldenbourg, 1918.

34. Pierrot-Deseilligny E, and Mazevet D. The monosynaptic reflex: a tool to investigate motor control in humans. Interest and limits. Neurophysiol Clin 30: 67–80, 2000.

35. Zehr EP, Collins DF, and Chua R. Human interlimb reflexes evoked by electrical stimulation of cutaneous nerves innervating the hand and foot. Exp Brain Res 140: 495–504, 2001.

36. Pearcey GEP, and Zehr EP. Repeated and patterned stimulation of cutaneous reflex pathways amplifies spinal cord excitability. J Neurophysiol 124: 342–351, 2020.

37. Hurteau MF, Thibaudier Y, Dambreville C, Danner SM, Rybak IA, and Frigon A. Intralimb and Interlimb Cutaneous Reflexes during Locomotion in the Intact Cat. J Neurosci 38: 4104–4122, 2018.

38. Desmedt HE, and Godaux E. Spinal motoneuron recruitment in man: rank deordering with direction but not with speed of voluntary movement. Science 214: 933–936, 1981.

39. ter Haar Romeny BM, van der Gon JJ, and Gielen CC. Relation between location of a motor unit in the human biceps brachii and its critical firing levels for different tasks. Experimental neurology 85: 631–650, 1984.

40. Keen DA, and Fuglevand AJ. Common input to motor neurons innervating the same and different compartments of the human extensor digitorum muscle. J Neurophysiol 91: 57–62, 2004.

41. McIsaac TL, and Fuglevand AJ. Motor-unit synchrony within and across compartments of the human flexor digitorum superficialis. J Neurophysiol 97: 550–556, 2007.

42. Rodriguez-Falces J, Negro F, and Farina D. Correlation between discharge timings of pairs of motor units reveals the presence but not the proportion of common synaptic input to motor neurons. J Neurophysiol 117: 1749–1760, 2017.

43. de la Rocha J, Doiron B, Shea-Brown E, Josic K, and Reyes A. Correlation between neural spike trains increases with firing rate. Nature 448: 802–806, 2007.

44. Christakos CN, Papadimitriou NA, and Erimaki S. Parallel neuronal mechanisms underlying physiological force tremor in steady muscle contractions of humans. J Neurophysiol 95: 53–66, 2006.

45. Laine CM, Yavuz SU, and Farina D. Task-related changes in sensorimotor integration influence the common synaptic input to motor neurones. Acta Physiol (Oxf 211: 229–239, 2014.

46. Glover IS, and Baker SN. Both Corticospinal and Reticulospinal Tracts Control Force of Contraction. J Neurosci 2022.

47. Cheung VC, d’Avella A, Tresch MC, and Bizzi E. Central and sensory contributions to the activation and organization of muscle synergies during natural motor behaviors. J Neurosci 25: 6419–6434, 2005.

48. Alessandro C, Barroso FO, Prashara A, Tentler DP, Yeh HY, and Tresch MC. Coordination amongst quadriceps muscles suggests neural regulation of internal joint stresses, not simplification of task performance. Proc Natl Acad Sci U S A 201916578, 2020.

49. Mang CS, Clair JM, and Collins DF. Neuromuscular electrical stimulation has a global effect on corticospinal excitability for leg muscles and a focused effect for hand muscles. Exp Brain Res 209: 355–363, 2011.

50. Pahwa R, Dhall R, Ostrem J, Gwinn R, Lyons K, Ro S, Dietiker C, Luthra N, Chidester P, Hamner S, Ross E, and Delp S. An Acute Randomized Controlled Trial of Noninvasive Peripheral Nerve Stimulation in Essential Tremor. Neuromodulation: journal of the International Neuromodulation Society 22: 537–545, 2019.

